# High-throughput Activity Assay for Screening Inhibitors of the SARS-CoV-2 Mac1 Macrodomain

**DOI:** 10.1101/2021.10.07.463234

**Authors:** Morgan Dasovich, Junlin Zhuo, Jack A. Goodman, Ajit Thomas, Robert Lyle McPherson, Aravinth Kumar Jayabalan, Veronica F. Busa, Shang-Jung Cheng, Brennan A. Murphy, Karli R. Redinger, Takashi Tsukamoto, Barbara Slusher, Jürgen Bosch, Huijun Wei, Anthony K. L. Leung

## Abstract

Macrodomains are a class of conserved ADP-ribosylhydrolases expressed by viruses of pandemic concern, including coronaviruses and alphaviruses. Viral macrodomains are critical for replication and virus-induced pathogenesis; therefore, these enzymes are a promising target for antiviral therapy. However, no potent or selective viral macrodomain inhibitors currently exist, in part due to the lack of a high-throughput assay for this class of enzymes. Here, we developed a high-throughput ADP-ribosylhydrolase assay using the SARS-CoV-2 macrodomain Mac1. We performed a pilot screen which identified dasatinib and dihydralazine as ADP-ribosylhydrolase inhibitors. Importantly, dasatinib does not inhibit MacroD2, the closest Mac1 homolog in humans. Our study demonstrates the feasibility of identifying selective inhibitors based on ADP-ribosylhydrolase activity, paving the way for screening large compound libraries to identify improved macrodomain inhibitors and explore their potential as antiviral therapies for SARS-CoV-2 and future viral threats.

---

Seven human coronaviruses have been identified: HCoV-229E, HCoV-NL63, HCoV-OC43 and HCoV-HKU1 are responsible for annual bouts of common cold while SARS-CoV, SARS-CoV-2, and MERS-CoV can cause severe pneumonia and are a major public health concern. Hundreds of additional coronaviruses are circulating in animal reservoirs and could be transmitted to humans.^1^ The diseases that result from zoonotic transfer are unpredictable, but historically are severe, highly contagious, and have potentially devastating consequences for public health. Therefore, developing broad-spectrum therapeutics against coronaviruses is of timely importance and will prepare us for future epidemics.

The SARS-CoV-2 genome encodes four structural proteins, nine accessory proteins, and 16 nonstructural proteins that are responsible for virus replication. COVID-19 antiviral development has focused on repurposing existing drugs to inhibit the enzymatic activities of proteins involved in SARS-CoV-2 replication, including viral RNA polymerases and proteases.^2^ As was the case for HIV and Hepatitis C virus, an effective treatment for SARS-CoV-2 will likely require a combination of drugs to pre-empt possible drug resistance. Therefore, identifying mechanistically distinct targets will complement current drug development efforts. Here we focus on screening for inhibitors of Mac1, a conserved macrodomain ADP-ribosylhydrolase within nonstructural protein 3 (nsp3).

Macrodomain is a protein fold found in humans and pathogens.^3–5^ Nearly all of them bind to adenosine diphosphate ribose (ADP-ribose).^4–7^ Recent data revealed that a subset of macrodomains hydrolyzes protein-conjugated ADP-ribose.^8–13^ For example, SARS-CoV, SARS-CoV-2 and MERS-CoV contain two to three macrodomains in tandem, where only the first one (called Mac1) possesses ADP-ribosylhydrolase activity.^13–17^ Notably, key residues critical for ADP-ribosylhydrolase activity are 100% conserved in all seven human coronaviruses as well as those identified from animal reservoirs, such as bat (Fig. S1). Macrodomain ADP-ribosylhydrolases are also conserved in another genus of pathogenic RNA viruses called alphaviruses (e.g. Chikungunya virus).^11,12^ Genetic evidence demonstrates the ADP-ribosylhydrolase activity of viral macrodomains is critical for replication and virulence.^11,13,18–21^ Mutant coronaviruses and alphaviruses cannot replicate when the ADP-ribose-binding sites within their macrodomains are disrupted.^11,21^ Additionally, macrodomain mutant viruses exhibit attenuated replication in differentiated cells and decreased virulence *in vivo*.^4,5^ Therefore, drugs targeting the ADP-ribosylhydrolase activity of viral macrodomains have the potential to inhibit viral replication and pathogenesis.

Two major challenges must be addressed during the development of antiviral macrodomain inhibitors. First, measurements of macrodomain ADP-ribosylhydrolase activity have historically relied on gel-based autoradiography and western blot assays that are not practical for screening large numbers of compounds. Second, humans express 11 proteins with macrodomain folds, such as MacroD2, which is the closest enzymatically-active human homolog of SARS-CoV-2 Mac1.^15^ Therefore, compounds that non-specifically inhibit human macrodomains will likely have off-target effects that limit their utility. Here we describe a quantitative, high-throughput assay that identified virus-specific and general inhibitors of macrodomains.

## Results and Discussion

To explore whether selective inhibition of a viral macrodomain is possible, we began our investigation by identifying biochemical and structural differences between SARS-CoV-2 Mac1 and human MacroD2. Sequence analyses classified both Mac1 and MacroD2 macrodomains to the macroD-type subclass, which also includes the macrodomain from the Chikungunya virus.^3,5^ Given that the Chikungunya virus macrodomain hydrolyzes ADP-ribose from recombinant PARP10 catalytic domain^11,12^ and G3BP1 protein from cells,^22^ we tested these substrates with Mac1 and MacroD2 (Fig. 1A and S2A). Following macrodomain incubation, comparable losses of ADP-ribose signal from PARP10^CD^ and G3BP1 were observed, indicating both SARS-CoV-2 Mac1 and human MacroD2 are active ADP-ribosylhydrolases.

**Fig. 1.**
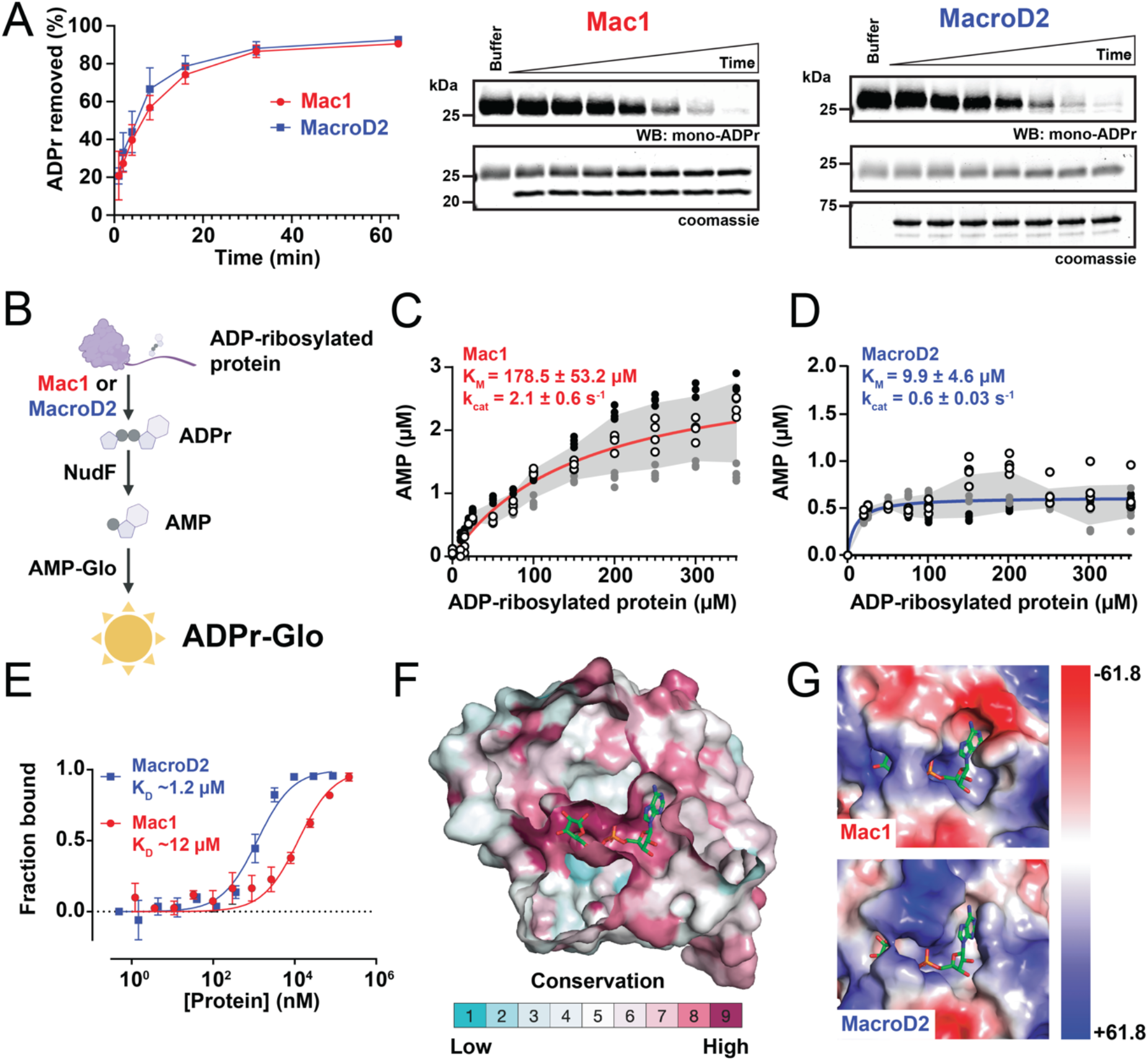
Biochemical, enzymatic, and structural characterization of SARS-CoV-2 Mac1 and human MacroD2. (**A**) Gel-based ADP-ribosylhydrolase assay against a mono-ADP-ribosylated substrate. Mono-ADP-ribose signal was normalized to the buffer signal. Plotted values are mean ± S.D. (n = 3). (**B)**Schematic of the luminescence-based ADP-ribosylhydrolase assay, ADPr-Glo. (**C-D**) Michaelis-Menten kinetics characterization of (C) Mac1 and (D) MacroD2 (n = 12, four technical replicates from three experiments, gray area is S.D.). (**E**) Electrophoretic mobility shift assay (EMSA) analyses of Mac1 and MacroD2 binding on Cy5-PAR. Plotted values are mean ± S.D., n = 3. (**F**) Surface representation of the conservation between Mac1 and MacroD2. Bound ADP-ribose is shown as stick representation. (**G**) Zoom-in view of the electrostatic surface potential of the ADP-ribose binding site for Mac1 (top) and MacroD2 (bottom).

To quantitatively measure the enzymatic activity with a high-throughput method, we developed the luminescence-based assay ADPr-Glo (Fig. 1B): First, ADP-ribose is released from a defined protein substrate by the macrodomain ADP-ribosylhydrolase. Second, the phosphodiesterase NudF cleaves the released ADP-ribose into phosphoribose and AMP. Finally, AMP is converted to luminescence with the commercially available AMP-Glo kit. This method takes advantage of the substrate selectivity of NudF, which cleaves free ADP-ribose but has no activity with protein-conjugated ADP-ribose.^23^ Therefore, the luminescence signal is controlled by the rate of the ADP-ribosylhydrolase. ADPr-Glo can be performed in 384-well plates with reaction volumes as low as 5 μL, greatly minimizing time and costs compared to gel-based activity assays.

We first used ADPr-Glo to measure the Michaelis-Menten kinetics of SARS-CoV-2 Mac1 and human MacroD2 with an ADP-ribosylated protein substrate. The K_M_ of Mac1 was 178.5 ± 53.2 μM with a k_cat_ of 2.1 ± 0.6 sec^−1^ (Fig. 1C), and the K_M_ of MacroD2 was 9.9 ± 4.6 μM with a k_cat_ of 0.6 ± 0.03 sec^− 1^ (Fig. 1D). The lower K_M_ of MacroD2 is consistent with its higher affinity for ADP-ribose monomers^16^ and polymers (Fig. 1E and S2B). Therefore, SARS-CoV-2 Mac1 and human MacroD2 exhibit distinct binding and kinetic properties with free and protein-conjugated ADP-ribose, which likely reflect chemical and structural differences within their active sites.

Comparison of the published macrodomain structures of Mac1 and MacroD2 revealed that ~60% of residues at the ADP-ribose binding sites are conserved (Fig. 1F). Similar to MacroD2 and the Chikungunya virus macrodomain,^9,11^ mutation of a conserved glycine residue to glutamate (G252E for SARS-CoV-2 nsp3, Fig. S1) abrogates the activity of Mac1 (Fig. S2C). Incubation of Mac1 G252E with the ADP-ribosylated protein substrate followed by NudF addition yielded the same amount of signal as the NudF-only control. A closer examination of Mac1 and MacroD2 structures revealed less conserved regions (e.g., the adenosine binding pocket; Fig. 1F) and distinctive electrostatic surfaces surrounding the active site where ADP-ribose binds (Fig. S2D). Compared with MacroD2, Mac1 possesses a binding pocket with more charged surfaces that is 450 Å^3^ larger (Fig. 1G and S2D-E). Taken together, these functional and structural differences may permit selective inhibition of SARS-CoV-2 Mac1, but not human MacroD2.

We next established ADPr-Glo conditions for inhibitor screening (Fig. 2A). The reaction was linear with respect to enzyme concentration (0.5 nM) and time of incubation (60 min) at room temperature in the presence of 20 μM ADP-ribosylated substrates, 125 nM NudF and final DMSO concentration of 1%, with excellent reproducibility when performed over different dates (Fig. S3). We then carried out a pilot screen of the 3,233 pharmacologically active compounds derived from the Selleck-FDA library (1,953) and the LOPAC library (1,280).

**Fig. 2.**
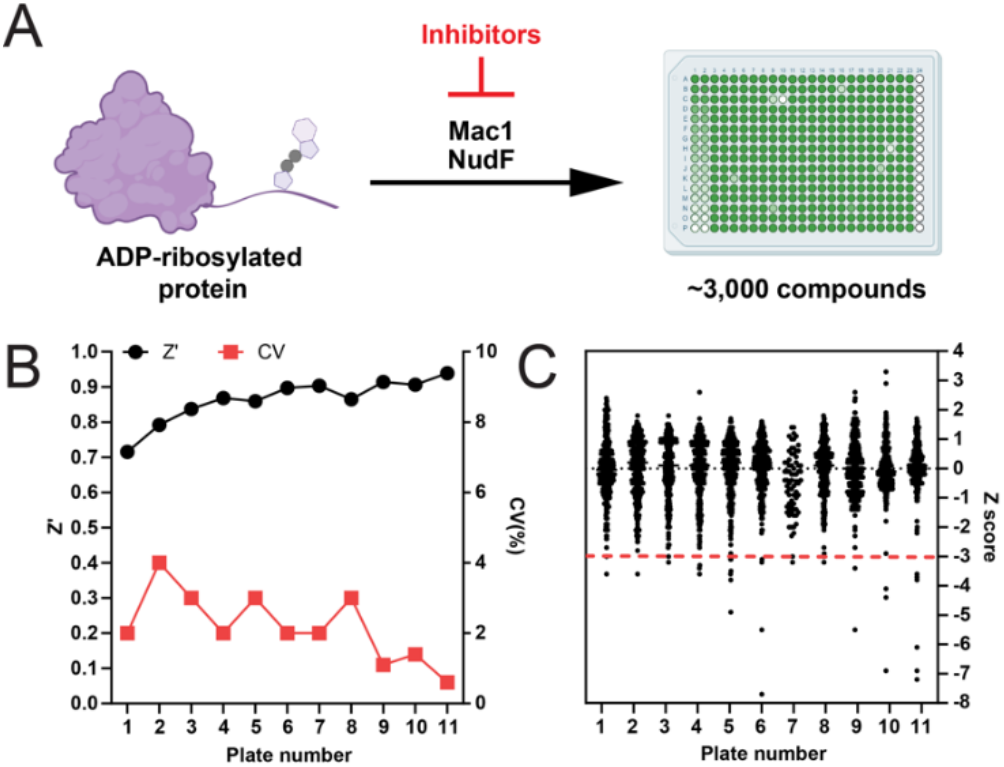
Pilot screen for macrodomain inhibitors. (A) Schematic of the drug screen based on the ADP-ribosylhydrolase assay ADPr-Glo. (B) Coefficients of variation (CV) and Z’ values for each plate in the screen. (C) Z scores for the 3,233 compounds evaluated.

The pilot screen parameters were suitable for a large high-throughput screen with the coefficient of variation (CV) ranging from 1-4%, the screening window coefficient Z’ at 0.86, and an average signal-to-background (S/B) ratio of 3.4 (Fig. 2B). We calculated the average signal (A) and standard deviation (SD) of compound-treated wells in each plate and determined a Z score for each compound where Z=(signal-A)/SD (Fig. 2C). Compounds with Z score ≤ −3 were considered hits (Supplementary Datafile 1).

Our pilot screen at 100 μM identified 21 compounds from the Selleck-FDA library and 16 compounds from the LOPAC library with Z ≤ −3, resulting in a 1.2% hit rate. Notably, the kinase inhibitor dasatinib was present as three different forms in the FDA library (Supplementary Datafile 1), and all of them were identified as hits, indicating assay reproducibility. Among 37 total hits, 24 were excluded based on several criteria, including the presence of pan-assay interfering (PAINS) substructures and/or potential aggregators based on the ZINC filtering algorithm^24^, interference of luminescence detection, high-molecular weight, instrument issues or commercial availability (see Supplementary Data-file 1). The remaining 13 hits were either purchased in powder form or synthesized for further evaluation.

To identify false-positive hits that either inhibit NudF or interfere with AMP detection by AMP-Glo, we performed a counter screen where 2 μM ADP-ribose was used instead of the ADP-ribosylated substrate and the macrodomain was omitted from the reaction (Fig. S4A-F). Four compounds demonstrated dose-dependent inhibition in the counter screen, indicating they are inhibitors of NudF and/or AMP-Glo (Fig. S4B-E). Vandetanib had poor solubility in aqueous solution (<10 μM) which prohibited dose-response analysis. The remaining six compounds did not inhibit the NudF-mediated counter screen assay and were subsequently evaluated in a dose-response assay against SARS-CoV-2 Mac1 (Fig. S4F-I).

Among the six remaining hits, only dasatinib and dihydralazine exhibited dose-dependent inhibition (Fig. 3A-B and S4G-I), with an IC_50_ of 37.5–57.5 μM and 485–757 μM (95% C.I.), respectively. We then evaluated these two inhibitors in an orthogonal gel-based activity assay. Consistent with its higher potency, only dasatinib mitigated the reduction of ADP-ribosylation under the tested conditions (Fig. 3C).

**Fig. 3.**
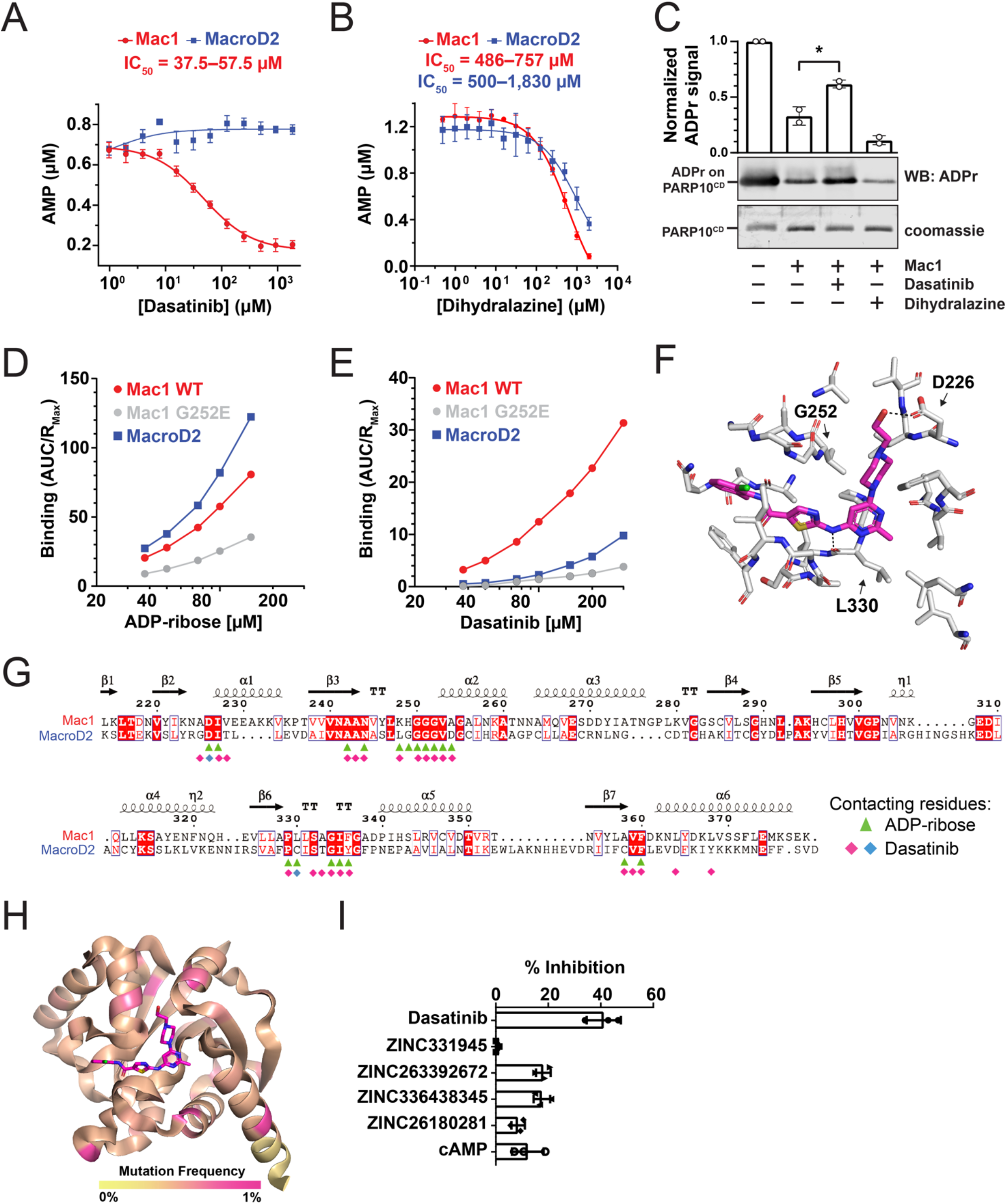
Dasatinib inhibits SARS-CoV-2 Mac1, but not human MacroD2. (**A-B**) Dose-response curves for (A) dasatinib and (B) dihydralazine against SARS-CoV-2 Mac1 and MacroD2. Plotted values are mean ± S.D. (n = 4). (**C**) Gel-based assay demonstrating inhibition of Mac1 by dasatinib (n = 2). (**D-E**) SPR analyses of (D) ADP-ribose and (E) dasatinib binding to Mac1 wild-type (WT) and G252E as well as MacroD2. The binding was quantitated by the Area Under Curve (AUC) normalized by the maximal response unit (R_max_). (**F**) Molecular docking of dasatinib to Mac1. (**G**) Structure-based sequence alignment of Mac1 and MacroD2 with contacting residues to ADP-ribose (green) and dasatinib, where red and blue indicates hydrophobic and hydrogen-bond (e.g., Asp 226 and Leu 330) interactions, respectively. (**H**) Analyses of 440,212 SARS-CoV-2 genomes revealed the dasatinib docking site is highly conserved. No residues within 5Å near the docking site have high mutation frequencies. (**I**) Comparison of dasatinib with other macrodomain inhibitor hits using ADPr-Glo with 100 μM inhibitor.

To evaluate whether these drugs broadly inhibit ADP-ribosylhydrolases or are specific for Mac1, we replaced the SARS-CoV-2 macrodomain with human MacroD2 in our ADPr-Glo assay and tested for dose-dependent inhibition (Fig. 3A-B). Strikingly, dasatinib did not show any inhibition of MacroD2 even at 2 mM, the solubility limit of the compound in 2% DMSO. On the contrary, dihydralazine inhibited MacroD2 and Mac1 with comparable potency (IC_50_ = 500-1,830 μM, *P* = 0.82, t-test).

Because dasatinib was more potent and selective, we focused our efforts on investigating the dasatinib–macrodomain interaction. To directly assess the dasatinib binding to these two macrodomains, we performed Surface Plasmon Resonance (SPR) analyses. Both Mac1 and MacroD2 bound strongly to ADP-ribose (Fig. 3D and S5), indicating that both macrodomains were properly folded and able to interact with small-molecule ligands in our SPR assays. Consistent with the selective inhibition observed in the ADPr-Glo assay, dasatinib bound ~3-fold more to Mac1 than MacroD2 (Fig. 3E and S5). Molecular docking analyses revealed that dasatinib binds at the highly conserved ADP-ribose binding site (Fig. 3F-H and S6, Supporting Datafile 2), which is supported by the lack of ADP-ribose and dasatinib binding by the active site mutant G252E (Fig. 3D-F and S5). Notably, ten of 25 dasatinib-contacting residues in Mac1 are not conserved in MacroD2 (Fig. 3G), which may explain the selectivity.

Recent high-throughput efforts have used virtual and binding screens to identify compounds and fragments that bind to SARS-CoV-2 Mac1.^25–27^ We directly compared dasatinib to the hits identified in these studies and found that dasatinib was a more potent ADP-ribosylhydrolase inhibitor (Fig. 3I and S7A) and a stronger Mac1 binder (Fig. S7B). Notably, dasatinib was not identified as a hit despite being included in libraries used by Schuller et al. One possibility is that dasatinib produced high fluorescence when mixed with SYPRO Orange, a dye commonly used in differential scanning fluorimetry (Fig. S7C) and may therefore be a false negative in prior screens. These findings collectively highlight the novelty and benefits of our functional screening approach as a complement to existing screens that assay binding.

In summary, we have established a new functional assay to identify ADP-ribosylhydrolase inhibitors. Our facile and versatile assay identifies both specific and general ADP-ribosylhydrolase inhibitors. Our pilot screen identified dasatinib, whose selectivity demonstrates it is possible to discover drugs that specifically inhibit viral macrodomains. Although cytotoxic when used at μM concentration,^28^ dasatinib has antiviral activities against SARS-CoV and MERS-CoV through an unknown mechanism.^29^ Therefore, data presented in this study provide strong support for our target and assay strategy, which can be applied to a large-scale high-throughput screen for new and improved viral macrodomain inhibitors. As the macrodomain fold is highly conserved in all coronaviruses and alphaviruses, this screening tool represents an important step towards developing new broad-spectrum antivirals.

## Supporting information

Pilot screen raw data

Conservation of SARS-CoV-2 genomes

## ASSOCIATED CONTENT

Materials and Methods; Supplementary Figures 1–7: Sequence alignment of coronavirus macrodomains, enzymatic and structural comparisons of SARS-CoV-2 Mac1 and MacroD2, optimizing assay parameters for drug screening, evaluation of pilot screen hits, surface plasmon resonance traces, molecular docking of dasatinib with SARS-CoV-2 Mac1, comparison of dasatinib with hits identified in previously published screens (PDF).

Supplementary Datafile 1: Pilot screen raw data (XLSX) Supplementary Datafile 2: Conservation of SARS-CoV-2 genomes (XLSX)

## AUTHOR INFORMATION

### Author Contributions

M.D., J.Z., J.A.G. and A.T. contributed equally to this work, and other contributions as follows: study conception: A.K.L.L.; research design: A.K.L.L., M.D., R.L.M., T.T., B.S., H.W; data collection: M.D., J.Z., J.A.G., A.T., A.K.J., K.R.R., J.B.; reagent generation: R.L.M., S-J.C., B.A.M; data analyses, V.F.B., K.R.R, J.B.; analysis and interpretation of results: M.D., J.B., H.W., A.K.L.L.; draft manuscript preparation: M.D. and A.K.L.L. All authors reviewed the results and approved the final version of the manuscript.

### Notes

The authors declare no competing financial interest.

## ACKNOWLEDGMENT

We thank Drs Diane Griffin, Mohsen Badiee, Rachy Abraham for their critiques of the manuscript, the NudF expression construct from Dr. Sandra Gabelli, Lauren Bambarger for the initial testing of the luminescence-based assay, and Dr. Said Goueli for advice on the AMP-Glo assay.

## ABBREVIATIONS

ADP: adenosine diphosphate
SARS: severe acute respiratory syndrome
MERS: middle east respiratory syndrome
CoV: coronavirus
HIV: human immunodeficiency virus

## Supporting Information

### Chemicals

Dasatinib (Cat# S1021), dasatinib Monohydrate (Cat# S7782), vandetanib (Cat# S1046), pixantrone maleate (Cat# S5059), allopurinol (Cat# S1630) and fludarabine phosphate (Cat# S1229) were purchased from SelleckChem.com, Tyrphostin AG 490 (Cat# T3434) from Sigma Aldrich, 2-methylthioadenosine diphosphate (Cat# 21230) from Cayman Chemical Company, Cyclic AMP (cAMP, Cat# 6099240) from PeproTech, SX048 (Compound ID: EN300-13489), SX051 (Compound ID: EN300-18713) and ZINC331945 (Compound ID: EN300-79874) from Enamine, ZINC336438345 (Cat# Z2093206487), ZINC263392672 (Cat# Z3977586993) and ZINC26180281 (Cat# Z1262625706) from MolPort. SCH-12679 was synthesized in the form of racemate as previously reported.^1^

### Protein multiple sequence alignment

FASTA files containing the macrodomain protein sequences were aligned with T-Coffee http://tcoffee.crg.cat/apps/tcoffee/do:regular.^2^ The .fasta_aln file downloaded from T-Coffee was then run in BoxShade https://embnet.vital-it.ch/software/BOX_form.html with the fraction of sequences that must agree for shading set to 0.5. The .rtf file downloaded from Boxshade was formatted in Adobe Illustrator. Conservation bar graphs were generated manually in Prism 9.0 (Graphpad) and formatted in Adobe Illustrator.

### Expression and purification of SARS-CoV-2 Mac1, MacroD2, PARP10 catalytic domain, and ADP-ribosylated protein substrates

All steps were performed on ice or at 4 °C unless otherwise noted. The SARS-CoV-2 macrodomain was expressed in E. coli BL21 DE3. The polynucleotide sequence of SARS-CoV-2 nsp3 (residues 200-380), i.e., Mac1, was cloned into a pSAT1 plasmid and fused with a 6xHis-tagged small ubiquitin-like modifier (HisSUMO) tag at its N-terminus to increase construct stability. Following transformation, a single colony was used to inoculate 50 mL of LB containing 100 μg/mL ampicillin and 1% w/v glucose, which was then grown at 37 °C, 200 rpm for 16-20 hr. 10 mL of the starter culture was used to inoculate 1 L of LB containing 100 μg/mL ampicillin, and the cells were grown at 37 °C, 200 rpm to an OD_600_ of ~0.8. At that point the incubator temperature was changed to 16 °C and the cells were grown for 1 h before IPTG was added to a final concentration of 0.5 mM. The protein was expressed at 16 °C, 200 rpm for 16-20 hr. Cells were harvested by centrifugation at 2,600 × g for 30 min, then cell pellets were stored at −20 °C until purification. The cell pellet was thawed and resuspended in 50 mL lysis buffer (20 mM sodium phosphate pH 7.5, 0.5 M NaCl, 50 mM imidazole, 10 mM 2-mercaptoethanol, 1% NP-40, 1X cOmplete EDTA-free protease inhibitor cocktail). The cells were lysed by sonication, then lysate was clarified by centrifugation at 24,000 × g for 30 min. The lysate was filtered, then stirred gently with HisPur Ni-NTA resin (2 mL) for 1 hr. The suspension was transferred to a polypropylene gravity column. The resin was washed with 20 mL wash buffer (20 mM sodium phosphate pH 7.5, 0.5 M NaCl, 50 mM imidazole, 10 mM 2-mercaptoethanol), then the protein was then eluted with 8 mL elution buffer (20 mM sodium phosphate pH 7.5, 0.5 M NaCl, 300 mM imidazole, 10 mM 2-mercaptoethanol). The eluted protein was concentrated to 2 mL and desalted with a HiTrap Desalting column (2 × 5 mL) on an NGC chromatography system into wash buffer. Concentration was estimated by NanoDrop with the equation [protein] = A280 / 11,920 M-1 cm-1. 6xHis-tagged SUMO endopeptidase (SENP) was added to a 50:1 HisSUMO-Mac1:SENP w/w ratio and the tube was incubated without shaking for 16 hr. HisPur resin (2 mL) was used to remove HisSUMO and SENP with the gravity column procedure described above. The flow through, containing untagged Mac1, was further purified by gel filtration on a Superdex 200pg HiLoad 16/60 equilibrated with gel filtration buffer (50 mM HEPES pH 7.5, 100 mM NaCl, 1 mM TCEP, 10% glycerol). Concentration was estimated by NanoDrop with the equation [protein] = A280 / 10,430 M-1 cm-1, then the protein was aliquoted, flash-frozen and stored at −80 °C. For protein used in SPR experiments, the tag cleavage step was omitted so the protein could be attached to the SPR chip via the 6xHis tag.

PARP10 catalytic domain and MacroD2 were purified as described (McPherson et al., 2017). For ADP-ribosylated substrate preparation, a plasmid encoding HisSUMO fused to the PARP10-derived sequence CRRPVEQVLYH was transformed in E. coli BL21 DE3 cells. A single colony was used to inoculate 50 mL of LB containing 100 μg/mL ampicillin and 1% w/v glucose, which was then grown at 37 °C, 200 rpm for 16-20 hr. 10 mL of the starter culture was used to inoculate 1 L of LB containing 100 μg/mL ampicillin, and the cells were grown at 37 °C, 200 rpm to an OD_600_ of ~0.8. The cells were then grown at 16 °C, 200 rpm for 1 h before IPTG was added to a final concentration of 0.5 mM. The protein was expressed at 16 °C, 200 rpm for 16-20 hr. Cells were harvested by centrifugation at 2,600 × g for 30 min, then cell pellets were stored at −20 °C until purification. The cell pellet was thawed and resuspended in 50 mL lysis buffer (20 mM sodium phosphate pH 7.5, 0.5 M NaCl, 50 mM imidazole, 10 mM 2-mercaptoethanol, 1% NP-40, 1X cOmplete EDTA-free protease inhibitor cocktail). The cells were lysed by sonication, then lysate was clarified by centrifugation at 24,000 × g for 30 min. The lysate was filtered, then stirred gently with HisPur Ni-NTA resin (2 mL) for 1 hr. The suspension was transferred to a polypropylene gravity column. The resin was washed with 20 mL wash buffer (20 mM sodium phosphate pH 7.5, 0.5 M NaCl, 50 mM imidazole, 10 mM 2-mercaptoethanol), then the protein was eluted with 8 mL elution buffer (20 mM sodium phosphate pH 7.5, 0.5 M NaCl, 300 mM imidazole, 10 mM 2-mercaptoethanol). The eluted protein was concentrated to 2 mL and further purified by gel filtration on a Superdex 200pg HiLoad 16/60 equilibrated with gel filtration buffer (50 mM HEPES pH 7.5, 100 mM NaCl, 1 mM TCEP, 10% glycerol. Concentration was estimated by NanoDrop with the equation [protein] = A_280_ / 2,980 M^−1^ cm^−1^, then the protein was aliquoted, flash-frozen and stored at −80 °C.

### Automodification of PARP10 protein

ADP-ribosylation of PARP10 catalytic domain was performed essentially as described by Alhammad et al. (2020). PARP10 (10 μM) was incubated for 20 min at 37° C with NAD+ (1 mM) in reaction buffer (50 mM HEPES pH 7.2, 150 mM NaCl, 0.2 mM DTT, 0.02% NP-40). Automodified PARP10 was then aliquoted and stored at −80 °C.

### Gel-based ADP-ribosylhydrolase assay

The SARS-CoV-2 and human macrodomains (1 μM) were incubated with ADP-ribosylated PARP10 catalytic domain (1 μM) at 37 °C for 1-64 min. The ADP-ribose on PARP10 was then detected by western blotting with an anti-mono-ADP-ribose binding reagent (Millipore MABE1076). Total protein levels were determined by the SimplyBlue stain (Invitrogen LC6065). Buffer reactions were incubated at 37 °C for 64 min in the absence of macrodomain. Mono-ADP-ribose signal was quantified in ImageStudio (Li-Cor) then normalized to the buffer signal from each respective experiment. Plotted values are mean ± S.D. (n = 3).

### Preparation of the ADP-ribosylated protein substrate

All steps were performed on ice or at 4 °C unless otherwise noted. PARP10 catalytic domain (1 μM), HisSUMO-CRRPVEQVLYH (20 μM) and NAD+ (600 μM) were combined in MARylation buffer (50 mM HEPES pH 7.2, 150 mM NaCl, 1 mM TCEP) in a reaction volume totaling 25 mL. The reaction was mixed by inversion, then incubated at ambient temperature for 16 hr. HisPur resin (2 mL) was added and the tube was incubated with gentle agitation for 30 min. The suspension was transferred to a polypropylene gravity column and the resin was washed with 50 mL wash buffer (50 mM MES pH 6.0, 0.5 M NaCl, 50 mM imidazole, 1 mM TCEP, 5% v/v glycerol) to remove PARP10. The remaining protein was eluted with 8 mL elution buffer (50 mM MES pH 6.0, 0.5 M NaCl, 300 mM imidazole, 1 mM TCEP, 5% v/v glycerol), then concentrated and buffer exchanged into storage buffer (50 mM MES pH 6.0, 0.5 M NaCl, 1 mM TCEP, 5% v/v glycerol) with an Amicon centrifugal filter (3,000 MWCO) by spinning at 4,000 × g in 15-min intervals. Concentration was estimated by NanoDrop with the equation [ADP-ribosylated protein] = A_260_ / 13,500 M^−1^ cm^−1^, then the protein was aliquoted, flash-frozen and stored at −80 °C.

#### Quantitative measurement of ADP-ribosylhydrolase activity with the luminescence-based assay, ADPr-Glo

ADPr is produced from hydrolysis of mono-ADP-ribosylated substrates, then free ADPr is cleaved into AMP and phosphoribose by the NudF phosphodiesterase. AMP is then converted to luminescence with the AMP-Glo kit (Promega V5012).

### Enzyme kinetic analyses

Mac1 (0.86 nM) and NudF (125 nM) were incubated with ADP-ribosylated substrate at ambient temperature for 30 min. Mac1 and NudF were removed with a Microcon spin filter (10,000 MWCO), then the reaction products were measured with AMP-Glo. Reactions without Mac1 were performed in parallel as a negative control. Luminescence signal was converted to AMP generated via interpolation from an AMP standard curve. Data plotted are AMP generated by Mac1 and NudF, subtracted by AMP generated from NudF alone (n = 12, four technical replicates from three independent experiments, with each day represented by a different color). Standard deviation from the mean is shown in gray. Kinetic parameters were calculated with a Michaelis-Menten non-linear regression in Prism 9.0 (GraphPad).

### Electrophoretic Mobility Shift Assay

The 20-mer Cy5-PAR was prepared as described by Abraham et al. (2020). Cy5-PAR (10 nM) was mixed with protein at the indicated concentration in binding buffer (10 mM Tris pH 7.5, 100 mM KCl, 1 mM EDTA, 0.1 mM DTT, 0.01 mg/mL BSA, 0.01% w/v OrangeG, 5% v/v glycerol) and incubated at ambient temperature for 1 h. Samples were separated with a native 5% trisacetate polyacrylamide gel (37.5:1 acrylamide:bis-acrylamide ratio) by applying 10 V per cm of gel in tris-acetate buffer (40 mM Tris, 2.5 mM EDTA, 20 mM acetate pH 7.8) for 45 min. Cy5 signal was detected with a Typhoon (Molecular Biosciences). Cy5 signal was quantified with ImageStudio (Li-Cor), then plotted in Prism 9.0 (GraphPad). Dissociation constants were calculated with a sigmoidal dose-response curve by measuring [protein] at which half of the Cy5-PAR is bound in Prism 9.0 (GraphPad).

### Orthogonal gel-based assay with ADP-ribosylhydrolase inhibitors

SARS-CoV-2 Mac1 (25 nM) was incubated with compound (400 μM, 2% DMSO) in reaction buffer (50 mM HEPES pH 7.2, 150 mM NaCl, 0.2 mM DTT, 0.02% NP-40) on ice for 30 min. ADP-ribosylated PARP10 (1 μM) was then added and the reaction was incubated for 1 h at 37° C. Reactions were quenched by addition of 3X lithium dodecyl sulfate sample buffer to a final concentration of 1X.

### Gel electrophoresis and western blotting

Demodification samples (13.33 pmol PARP10^CD^ per lane) were separated with SDS-PAGE using 4-12% BisTris gels in MOPS-Tris running buffer, then transferred onto PVDF membranes. After a 4 °C overnight incubation in blocking solution (5% w/v non-fat dry milk in TBS-T + 0.02% w/v sodium azide), membranes immunoblotted using the mono-ADP-ribose detection reagent (2 μg/mL in blocking solution, Millipore MABE1076) for 1 h at ambient temperature. Following three 5 min washes in TBS-T, membranes were immunoblotted with secondary anti-Rabbit IgG polyclonal antibody conjugated with IRdye 800 (1:10,000 in blocking solution, Li-Cor 926-32213) for 1 h at ambient temperature. Following three 5 min washes in TBS-T, membranes were visualized with an Odyssey and data were quantified in ImageStudio.

### Pilot screen

Beckman ECHO acoustic liquid handler was used to transfer 50 nL of DMSO (columns 1-2, 23-23) or compounds (columns 3-22) from both the FDA-approved drugs screening library and Sigma’s LOPAC®1280 library to Greiner’s 384-well luminescence plates (Cat# 784904; small volume, non-binding). The day before the experiment, the plates were moved from the −80°C freezer to a 4°C refrigerator and then allowed to warm up to room temperature prior to the experiment. To control for inter-plate variability, serial dilutions of AMP were made up in 1X AMP-Glo buffer (50 mM HEPES, pH 7.0, 150 mM NaCl, 1 mM MgCl_2_, 0.01% Triton X-100, 2 mM β-mercaptoethanol) and 5 μL dispensed to columns 1-2 using an electronic multichannel pipettor. To initiate the experiment, solutions of Mac1 (0.83 nM) and NudF (104 nM) or NudF alone (104 nM) were made up in 1.67X AMP-Glo buffer and 3 μL of each dispensed to columns 3-23 and column 24, respectively. After 30 min, 2 μL of the ADP-ribosylated substrate described above (50 μM), diluted to the appropriate concentration in ultrapure water, was added to columns 3-24 and the mixture incubated for an additional 30 min. Following this incubation, the AMP generated by the Mac1, in the presence of compounds and/or DMSO, was detected using Promega’s AMP-Glo™ assay kit per manufacturer’s instructions. Briefly, 5 μL of AMP-Glo™ Reagent I was added to wells, mixed gently and the plates incubated for 60 min. Before the end of this incubation, and immediately before use, AMP-Glo™ Reagent II was added to the Kinase-Glo® One solution (at a ratio of 1:100 v/v) and 10 μL of this mixture added to all the wells, mixed gently and the plates incubated for another 60 min protected from light. Ensuing this final incubation, the plates were read in a BMG CLARIOStar luminometer and the full light captured from each well for 1 s. All incubations were carried out at room temperature. Unless specified, all reagents were dispensed using Thermo Scientific’s Multidrop instruments. Also, following every reagent addition, the plates were spun down for 1 min at 150 × g.

### Confirmation of initial hits and counter screen

All hits were then counter-screened to preclude assay-interfering compounds. Drugs were diluted to desired concentration in 2% DMSO and pre-incubated with NudF (125 nM) at ambient temperature for 30 min. 2 μL ADP-ribose solution was added to all wells at a final concentration of 2 μM and incubated with drug and NudF for 30 min. Following the incubation, 5 μL of AMP-Glo™ Reagent I was added to wells, mixed gently and the plates incubated for 60 min. Before the end of this incubation, and immediately before use, AMP-Glo™ Reagent II was added to the Kinase-Glo® One solution (at a ratio of 1:100 v/v) and 10 μL of this mixture added to all wells, mixed gently and the plates incubated for another 60 min. The luminescence was detected by a Synergy™ H1 Microplate Reader. Compounds that passed the counter screen will be selected for activity confirmation where they were preincubated with NudF (125 nM) alone or together with either Mac1 (1.7 nM) or MacroD2 (2 nM) for 30 min and then reacted with ADP-ribosylated substrates (20 μM) for 30 min.

### SARS-CoV-2 genome analyses

The nsp3 SARS-CoV-2 nucleotide sequence (positions 2720-8554 of NCBI Reference Sequence NC_045512.2) was aligned to 449,612 full-length SARS-CoV-2 genomes from the COVID-19 Data Portal in R using the package Biostrings pairwiseAlignment function.^3^ The aligned sequences were translated and trimmed to amino acid positions 207-373 of the nsp3 protein, which corresponds to the Mac1 macrodomain as in 6Z5T. Poor-quality sequences, defined as having greater than five mismatched or missing amino acids within the 167-amino acid macrodomain, were removed. The remaining 440,212 high-quality macrodomain sequences were then used to calculate mutation frequencies. All code necessary to reproduce the analysis as well as processed intermediate data are available at https://github.com/vbusa1/nsp3_macrodomain.

### Structure based sequence alignment, analysis of conservation and electrostatic potential analyses

The crystal structures of Mac1 (65ZT) and MacroD2 (4IQY) were used to generate a structure-based sequence alignment using ESPRESSO a subroutine of T-Coffee.^4^ Residues in 5Å proximity of the ADPr binding site were defined as the ADPr-binding pocket and compared for identity. The conservation plot was generated using the Consurf server^5^ using default values and the Mac1 (6Z5T) as a template. Vaccum electrostatics potential were calculated using PyMol 2.3.4 (The PyMOL Molecular Graphics System, Version 2.3 Schrödinger, LLC.).

### Molecular Docking studies

The following receptors were utilized for docking 6Z5T and 4IQY as a representative for SARS-CoV-2 and human MacroD2. Dasatinib was docked using FRED with a conformer library generated with Omega2^5–7^ using a high resolution grid. Visualization and analysis were carried out in Vida v4.4.0 (OpenEye Scientific Software). While many of the residues lining the active site pocket are conserved between the two proteins, the electrostatic surface potential varies significantly as well as the pocket size. Importantly, the ribose moiety in the MacroD2 is located in a more neutrally charged region, while the SARS-CoV-2 region is strongly positively charged. Similar charge reversal differences were observed in the adenine binding region, where the SARS-CoV-2 region is negatively charged while the MacroD2 is positively charged. These charge differences are likely the reason for very weak interaction of dasatinib with MacroD2. Additionally, the SARS-CoV-2 pocket is larger with approximately 1300 Å^3^ in size while the Human MacroD2 pocket is only 850 Å^3^ large. While ADP-ribose in both structures is stabilized by multiple polar interactions, dasatinib only makes two polar contacts based on our lowest energy binding pose. The majority of the contacts are of hydrophobic nature while the hydroxyl of sidechain Asp226 and the backbone oxygen of Leu 330 are hydrogen bonding with dasatinib.

### Surface Plasmon Resonance

All interactions were performed on a Biacore T200 instrument at 25°C using an HBS-P buffer (10 mM Hepes, 150 mM NaCl, 3 μM EDTA, 0.05% Surfactant P20, pH 7.4) supplemented with 0.5% DMSO. Proteins were covalently coupled to a CM5 chip at high density > 30000 RUs. Analytes were passed over the flow cells at six twofold dilutions with 60s contact and 120s dissociation time at a 30 μL/min flow rate. After each injection the flow cells were regenerated with a pulse of 10 mM Glycine pH 9. Analysis was carried out with Scrubber2 (BioLogic Software, http://www.biologic.com.au/scrubber.html) using the double referencing method and correcting for DMSO differences in the analyte samples. Data was corrected and normalized for the differences in molecular weight of the ligands as well as the captured protein molecular weights. Data visualization was performed in Prism 9.0 (GraphPad).

### Differential Scanning Fluorimetry

Experiments were performed essentially as described by Virdi et al. (2020). Briefly, a master mix of MOPS pH 7.0 (20 mM), NaCl (25 mM), SYPRO Orange (6.25X) and Mac1 (5 μM) was prepared and 19 μL was dispensed into each well of a 96-well PCR plate. Compound (500 μM, 1 μL of a 10 mM stock, n = 3) was added, then the plate was inserted into a 7500 Fast RT-qPCR (Applied Biosystems). Samples were incubated with a temperature gradient from 25 °C to 95 °C for 180 min while monitoring fluorescence in the “TAMRA” channel. Data were exported as a .csv and visualized as mean ± S.D. in Prism 9.0 (GraphPad).

## SUPPLEMENTARY FIGURES

**Fig. S1.**
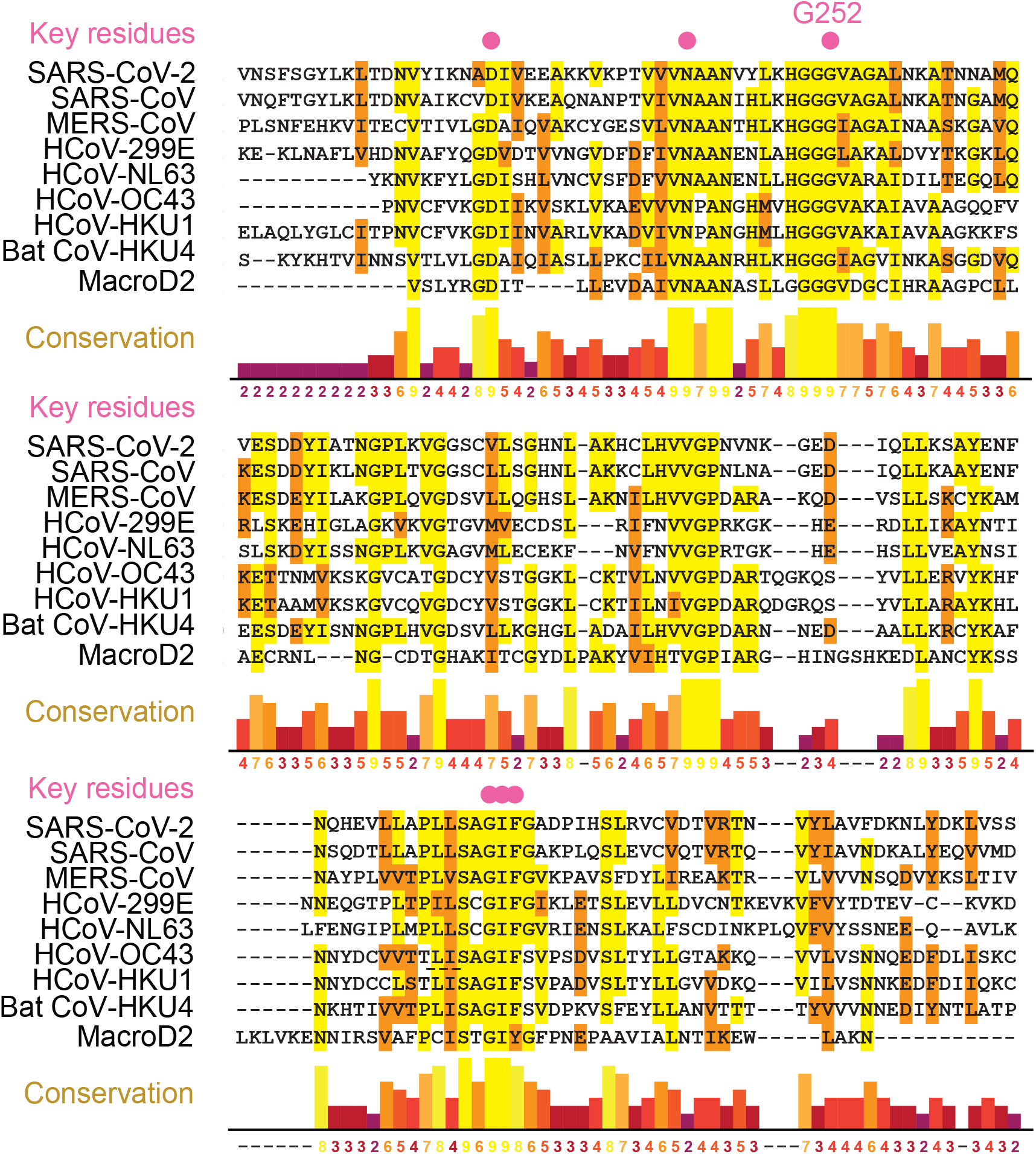
Multiple protein sequence alignment for seven human coronavirus macrodomains, one bat coronavirus macrodomain and the closest human homolog MacroD2. Red circles indicate residues that are critical for ADP-ribosylhydrolase activity based on previous studies (reviewed in Fehr *et al.*, 2018 and Leung *et al.*, 2018).

**Fig. S2.**
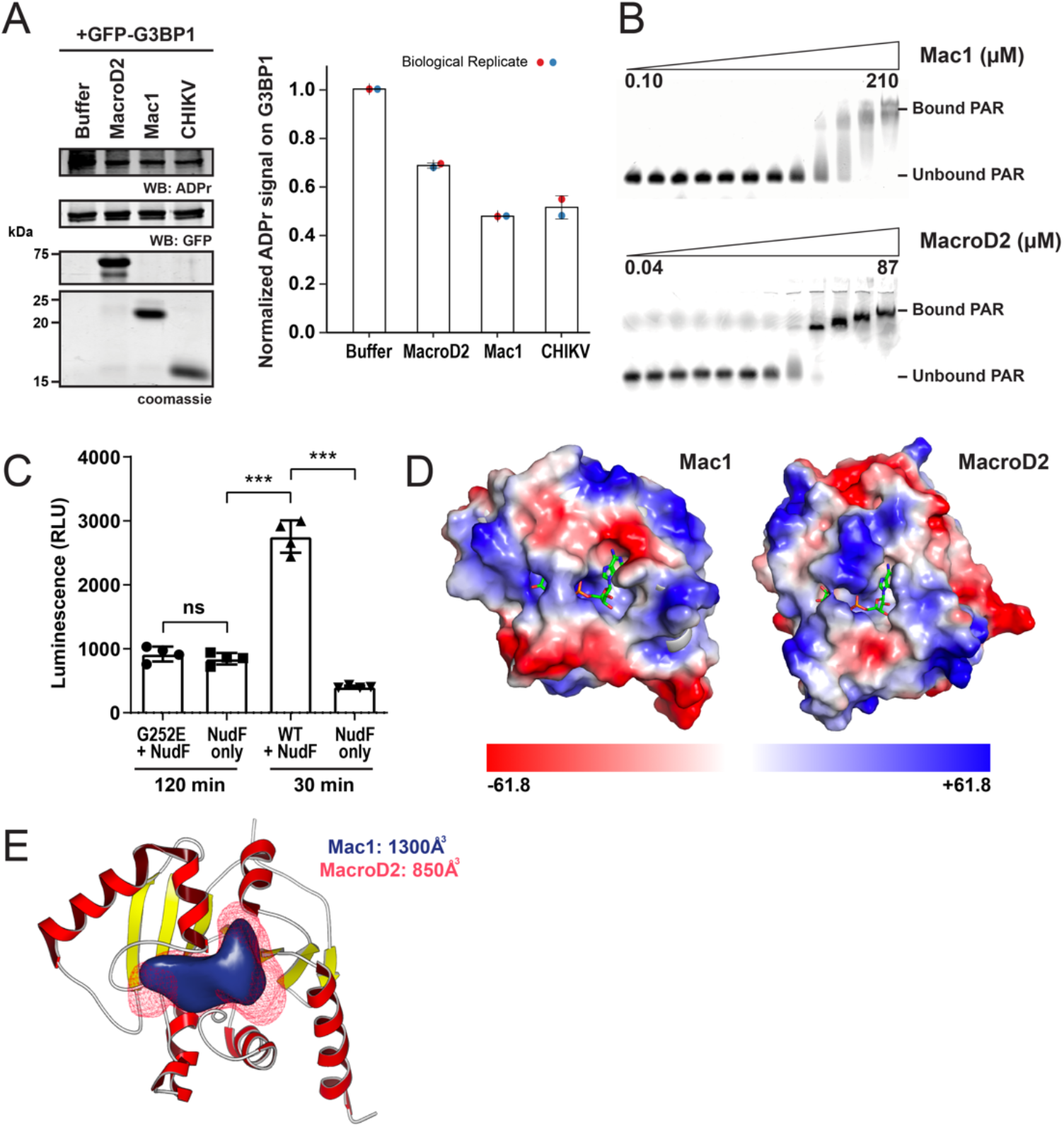
Biochemical, enzymatic, and structural characterization of SARS-CoV-2 Mac1 and human MacroD2. (**A**) Comparison of the ability of SARS-CoV-2 Mac1, MacroD2 and Chikungunya virus (CHIKV) macrodomain to remove ADP-ribose from ADP-ribosylated G3BP1 (Jayabalan et al., 2021). (**B**) Representative EMSA gels of Mac1 and MacroD2 binding to 20-mer Cy5-PAR. (**C**) The G252E mutation inactivates Mac1. Reactions with ADP-ribosylated substrate (20 μM) and wild-type Mac1 or G252E mutant (0.86 nM) were incubated at 37 °C for 30 min or 2 h, respectively. P-values were calculated with a 1-way ANOVA test; ***, p < 0.001. (**D**) Comparison of electrostatic surface potential between Mac1 and MacroD2. Red surfaces indicate negative charge and blue surfaces indicate positive charge. (**E**) Volume representation of the solvent accessible ADP-ribose binding pockets of Mac1 and MacroD2.

**Fig. S3.**
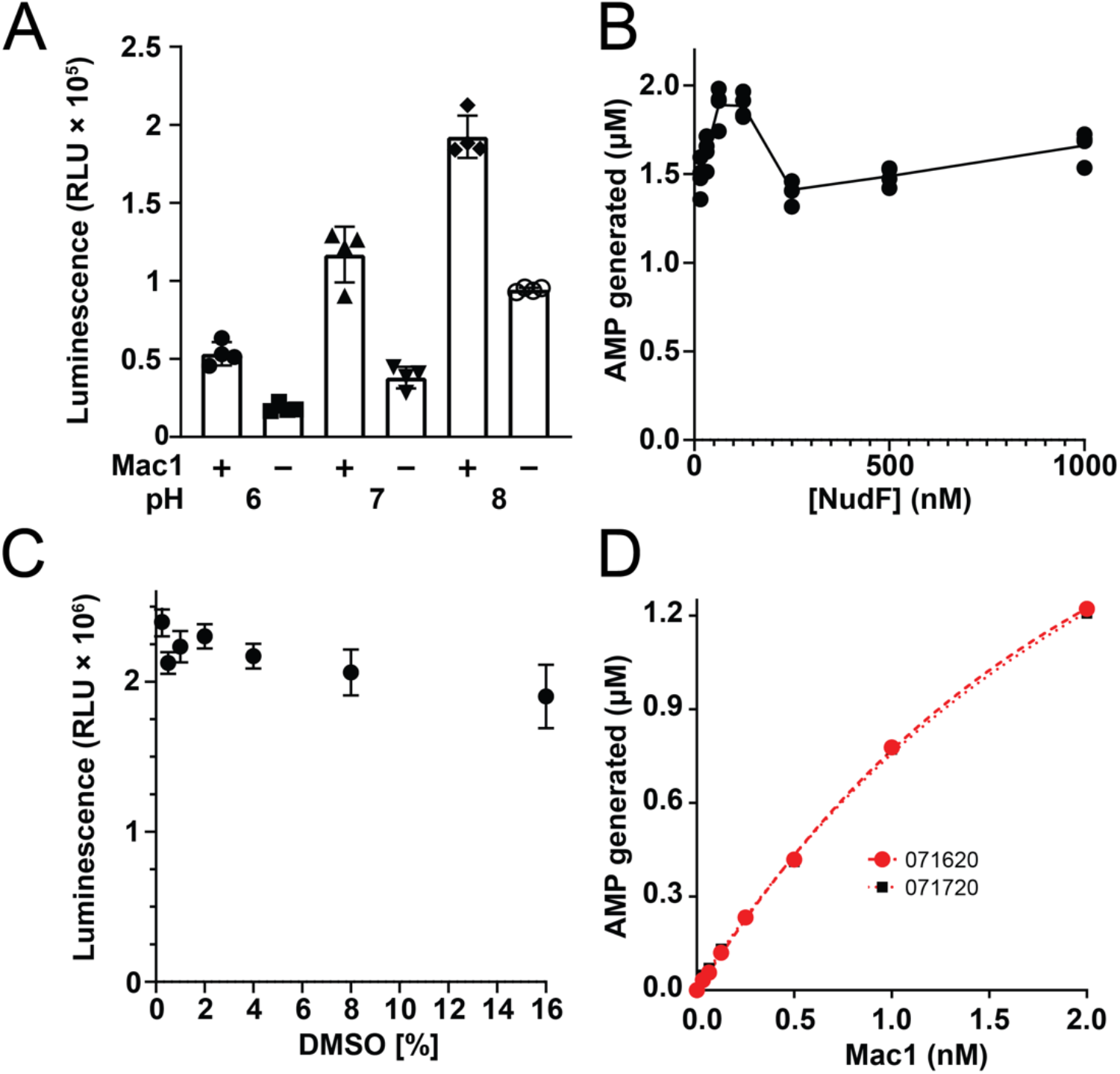
Determining optimal assay parameters for high-throughput screening. (**A**) pH dependence of luminescent signal from the ADPr-Glo assay. pH 7 has the highest signal-to-noise ratio. (**B**) AMP generated as a function of NudF concentration (15 nM – 1 μM). 125 nM was used for all kinetic assays and drug screening. Therefore, NudF is used in a large excess of the macrodomain such that the conversion of ADP-ribose to AMP is not the rate-limiting step. (**C**) The effect of DMSO concentration on the luminescent signal from the ADPr-Glo assay. (**D**) AMP generated as a function of SARS-CoV-2 macrodomain concentration (0.1 nM – 2 nM). The linear range is ≤ 1 nM, which was used for all kinetic assays and drug screening.

**Fig. S4.**
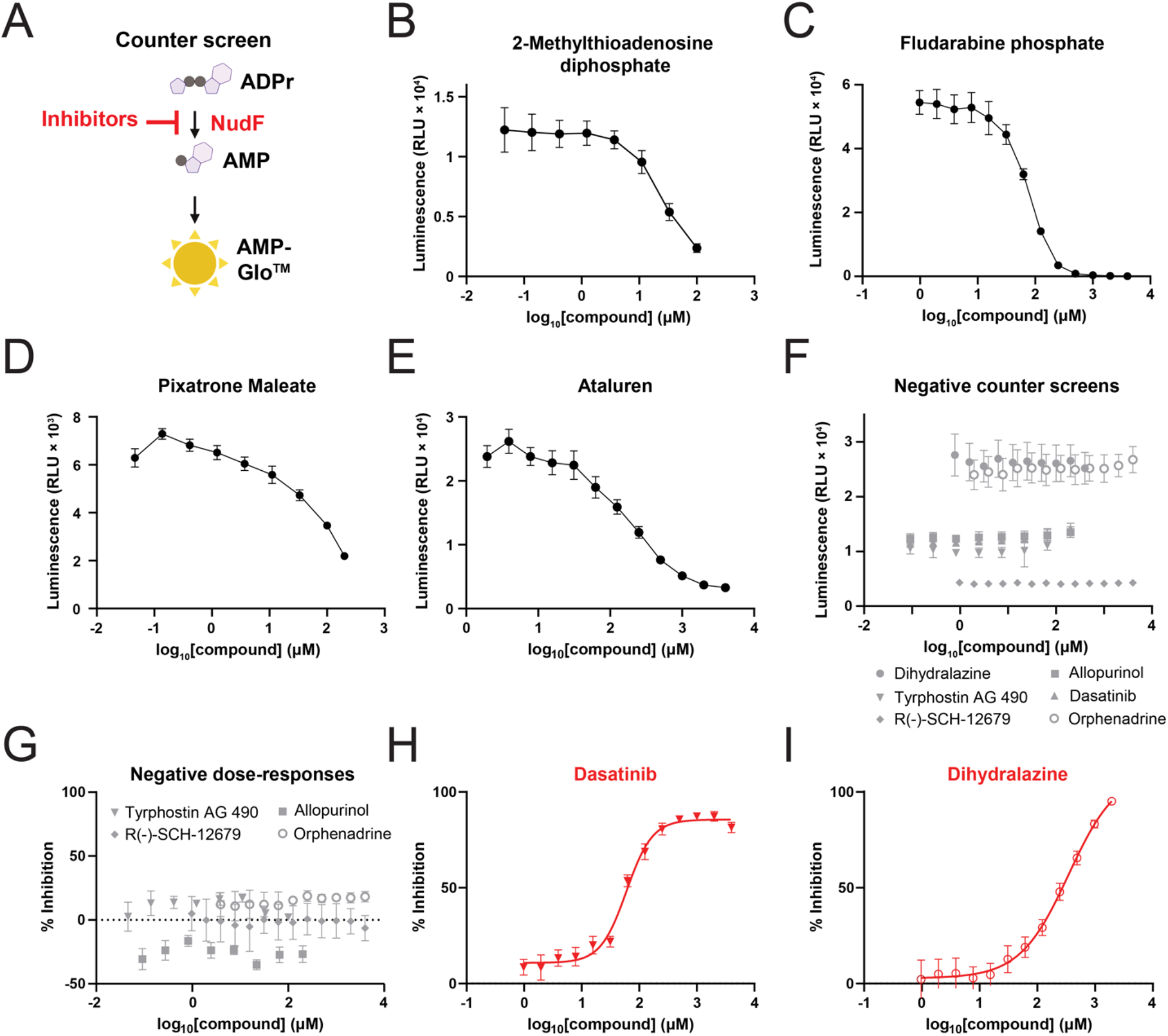
Evaluation of pilot screen hits. (**A**) Schematic for the counter-screening procedure. To identify compounds that reduce signal in the assay independent of the macrodomain, free ADPr (2 μM) replaces the ADP-ribosylated substrate and Mac1 is excluded. (**B-F**) Counter screens identified 2-methylthioadenosine diphosphate, fludarabine phosphate, pixatrone maleate, and ataluren as inhibitors of NudF and/or AMP-Glo. (**G-I**) Dose-response curves of the candidates that passed NudF/AMP-Glo counter screen using ADPr-Glo assay. Only dasatinib and dihydralazine demonstrated dose-dependent inhibition of SARS-CoV-2 Mac1.

**Fig. S5.**
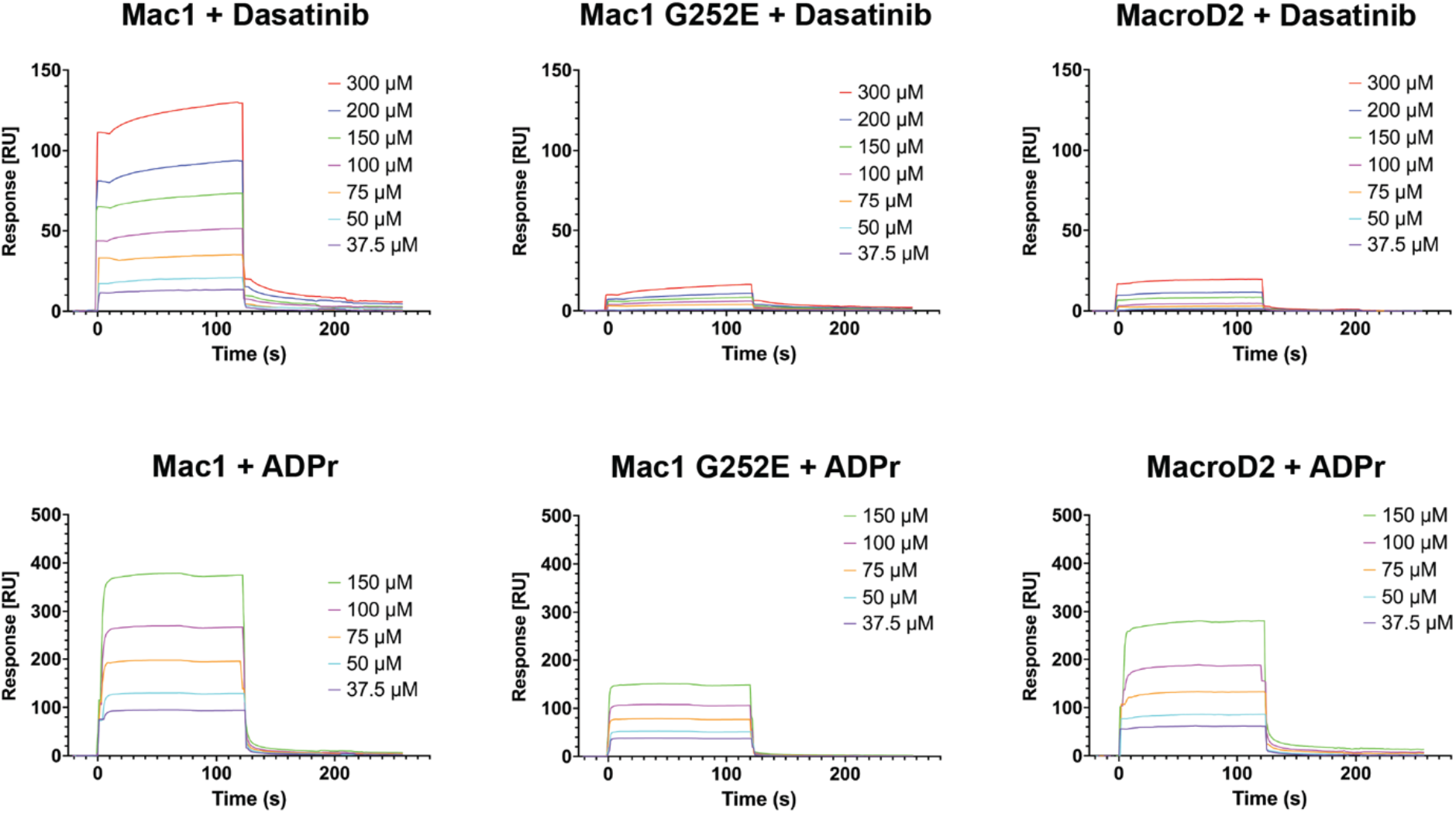
Surface Plasmon Resonance Analyses. The binding of (**A**) Dasatinib and (**B**) ADP-ribose (ADPr) with Mac1 WT and G252E as well as MacroD2.

**Fig. S6.**
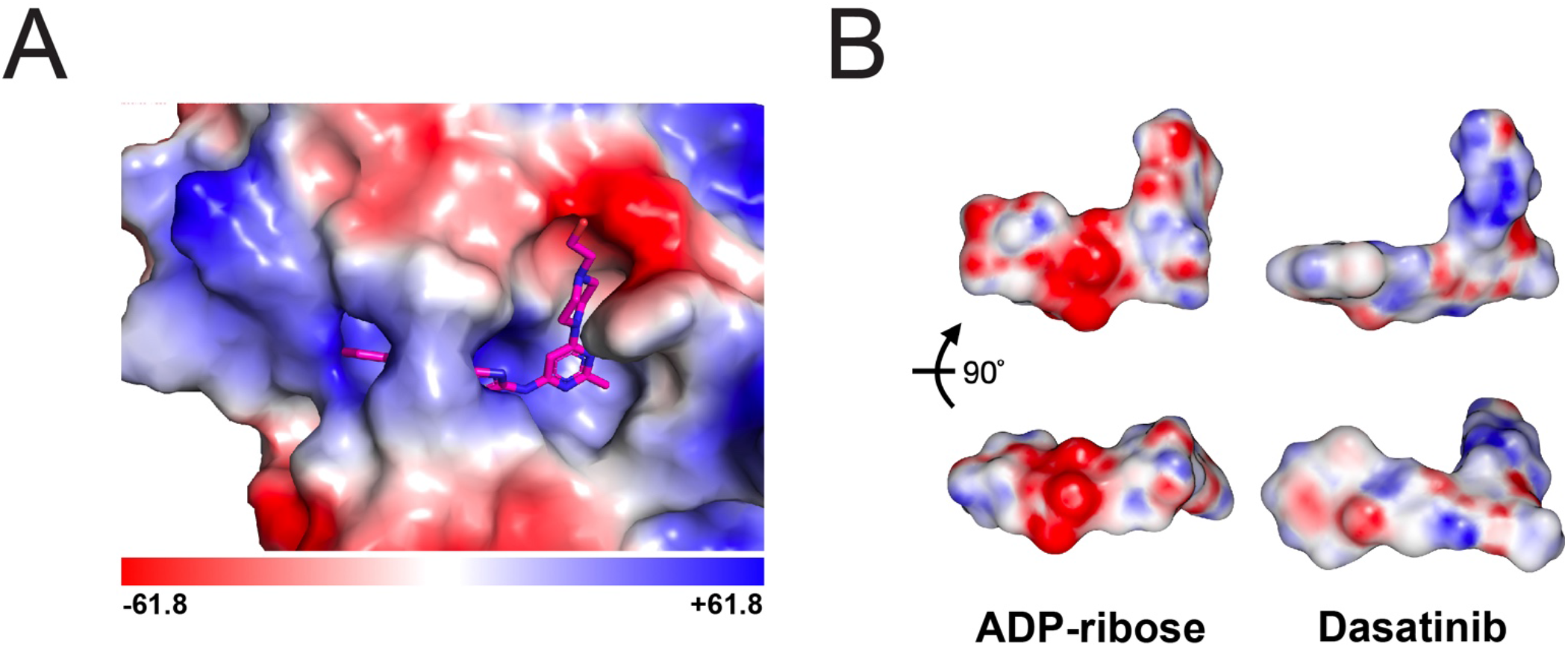
Molecular Docking of dasatinib with SARS-CoV-2 Mac1. (**A)**Electrostatic surface potential of the area surrounding the dasatinib docking site, with red indicating a negative potential and blue indicating a positive. (**B**) Electrostatic surface potential of ADP-ribose and dasatinib in two orientations. The overall shapes of the two molecules are comparable though distinct in their chemical properties. We note that the large negative ring of ADP-ribose snuggly interacts with the corresponding countercharge in the binding pocket, but dasatinib provides only a small negatively charged area in the corresponding location.

**Fig. S7.**
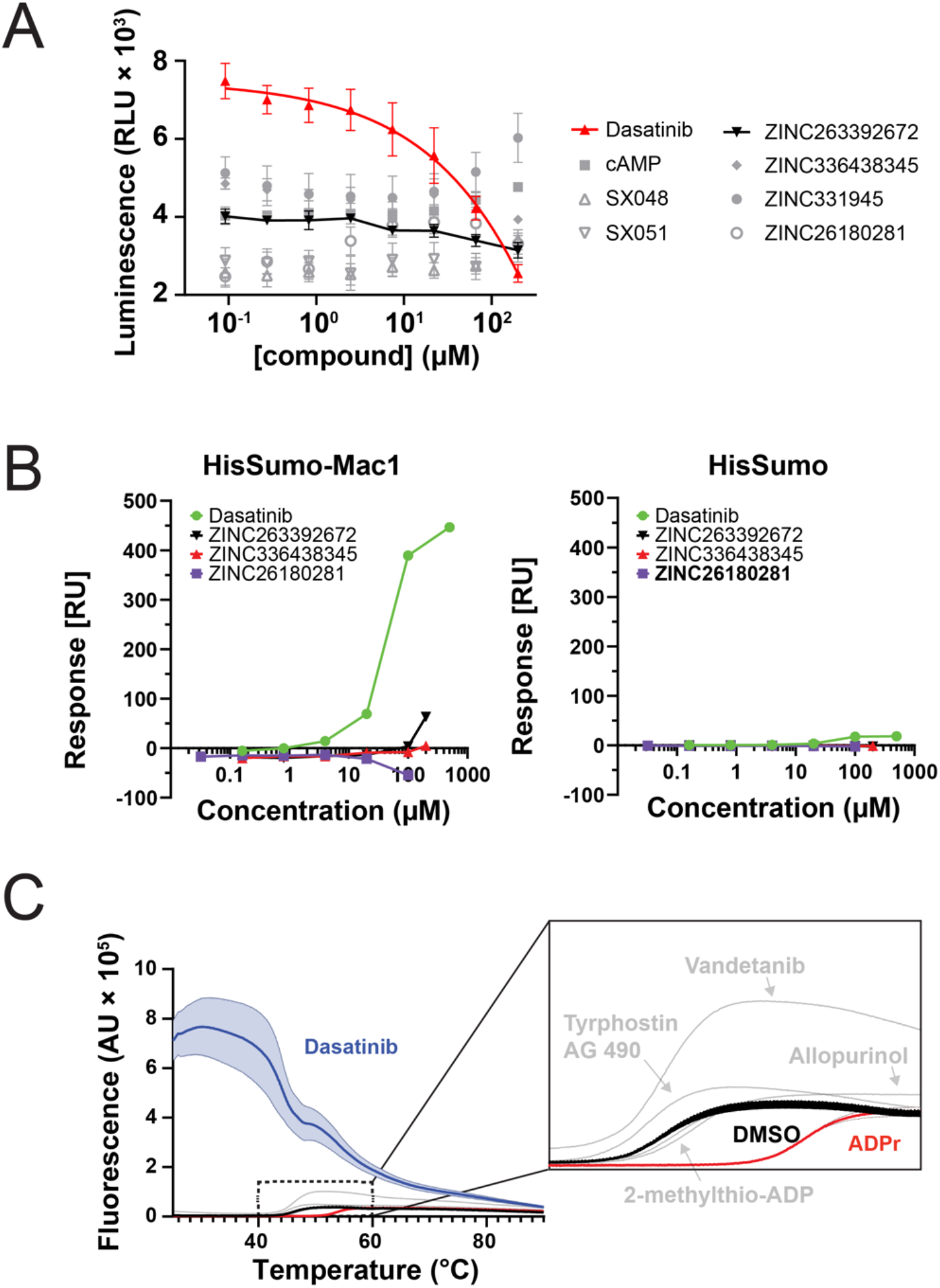
Comparison of dasatinib with other macrodomain inhibitor hits from published screens that are based on binding assays. (**A**) ADPr-Glo dose-dependent analyses of various compounds or fragments, including ZINC331945, trifluoroacetic acid (ZINC263392672), 3-fluoro-1-(7H-pyrrolo[2,3-d]pyrimidin-4-yl]piperidine (ZINC336438345), 9-methyl-2,6-diaminopurine (ZINC26180281) by Schuller et al., 2021, 1-Carbamoylpiperidine-4-carboxylic acid (SX048),(5S)-1-(4-Chlorophenyl)-5-methylimidazolidine-2,4-dione (SX051) from Bajusz *et al.*, 2021, and cAMP by Virdi *et al.*, 2020. (**B**) SPR analyses of dasatinib and fragment hits by Schuller et al with HisSumo-Mac1. cAMP, SX048, SX051, ZINC331945 were not analyzed because they did not inhibit Mac1 (panel A). While the fragments have been crystallized at very high concentrations, we did not observe a significant dose-dependent binding to Mac1 at the concentrations tested, except for ZINC263392672 at its highest concentration. Notably, ZINC263392672 also inhibited weakly Mac1 in a dose-dependent manner (panel A). Clear binding of dasatinib was observed to His-Sumo-Mac1 but not His-Sumo alone, indicating specific binding to SARS-CoV-2 Mac1. (**C**) Differential scanning fluorimetry of Mac1 with five hits from our screen, with ADP-ribose as a positive control and DMSO as a negative control. Dasatinib (blue) exhibits high background fluorescence and thus cannot be evaluated with this assay. Plotted values are mean ± S.D. (n = 3).

